# Quantifiable *In Vivo* Imaging Biomarkers of Retinal Regeneration by Photoreceptor Cell Transplantation

**DOI:** 10.1101/2019.12.21.884973

**Authors:** Ying V. Liu, Simrat Sodhi, Gilbert Xue, Derek Teng, Dzhalal Agakishiev, Minda M. McNally, Sarah Harris-Bookman, Caitlin McBride, Gregory Konar, Mandeep S. Singh

## Abstract

**Purpose:** Short-term improvements in retinal anatomy are known to occur in preclinical models of photoreceptor transplantation. However, correlative changes over the long term are poorly understood. We aimed to develop a quantifiable imaging biomarker grading scheme, using non-invasive multimodal confocal scanning laser ophthalmoscopy (cSLO) imaging, to enable serial evaluation of photoreceptor transplantation over the long term.

**Methods:** Yellow-green fluorescent microspheres were transplanted into the vitreous cavity and/or subretinal space of *C57/BL6J* mice. Photoreceptor cell suspensions or sheets from rhodopsin-green fluorescent protein mice were transplanted subretinally, into either *NOD.CB17-Prkdc^scid^/J* or *C3H/HeJ-Pde6b^rd1^* mice. Multimodal cSLO imaging was performed serially for up to three months after transplantation. Imaging biomarkers were scored, and a grade was defined for each eye by integrating the scores. Image grades were correlated with immunohistochemistry (IHC) data.

**Results:** Multimodal imaging enabled the extraction of quantitative imaging biomarkers including graft size, GFP intensity, graft length, on-target graft placement, intra-graft lamination, hemorrhage, retinal atrophy, and peri-retinal proliferation. Migration of transplanted material was observed. Changes in biomarker scores and grades were detected in 13/16 and 7/16 eyes, respectively. A high correlation was found between image grades and IHC parameters.

**Conclusions:** Serial evaluation of multiple imaging biomarkers, when integrated into a per-eye grading scheme, enabled comprehensive tracking of longitudinal changes in photoreceptor cell grafts over time. The application of systematic multimodal *in vivo* imaging could be useful in increasing the efficiency of preclinical retinal cell transplantation studies in rodents and other animal models.

## Introduction

Photoreceptor transplantation is being developed as a therapeutic modality to restore vision in people affected by retinal degenerative diseases, including retinitis pigmentosa (RP) and age-related macular degeneration (AMD) ^1–6^. Short term improvements in outer retinal anatomy after photoreceptor cell transplantation have been observed, mainly by histological staining in preclinical models of retinal degeneration ^4,7^. However, histology is a relatively inefficient method to track graft and recipient anatomy longitudinally over the long term. Histological assays are labor-intensive, require large initial cohorts of recipients, and face the challenge of recipient attrition over time. Non-invasive imaging could facilitate the longitudinal evaluation of retinal anatomy in relatively smaller cohorts of recipient animals over time, without the need to sacrifice animals at every assessment time point.

Recent advances in imaging techniques have enabled detailed imaging studies in mouse models of retinal disease and regeneration ^8–10^. Confocal scanning laser ophthalmoscopy (cSLO) can be used to capture images in multiple imaging modes, including short-wavelength fluorescence (SWF) excitation (488 nm) to detect photoreceptor cells labeled with green fluorescent protein (GFP) in mouse recipients ^1^. Multicolor reflectance (MR) imaging combines blue (488 nm), green (515 nm), and infrared (820 nm) laser reflectance. MR imaging has been applied to evaluate retinopathies such as geographic atrophy ^11–13^ and retinal edema ^13,14^ in humans, but has not yet been explored comprehensively as a tool in preclinical photoreceptor transplantation research. Conventional time-domain optical coherence tomography (OCT) has been applied to study transplantation outcomes in the short term ^15–18^. Spectral domain OCT (SD-OCT) produces high-resolution images, enabling the identification of distinct retinal layers in the recipient ^19–22^.

The utility of multimodal *in vivo* photoreceptor graft imaging for long-term observation has not yet been fully explored. We hypothesized that a non-invasive multimodal *in vivo* imaging system would provide comprehensive longitudinal information regarding graft survival and recipient retinal status. We aimed to develop quantifiable imaging biomarkers of photoreceptor transplantation outcomes in a mouse model based on multimodal *in vivo* cSLO imaging. We further aimed to develop a comprehensive imaging-based grading system to evaluate retinal changes in a cohort of transplant recipients over time.

## Methods & Materials

### Ethics approval

All animal experiments were carried out in accordance with the ARVO Statement for the Use of Animals in Ophthalmic and Vision Research. All procedures were approved by the Johns Hopkins University Animal Care and Use Committee (approval M016M17).

### Animals

Wild-type *C57/BL6J* mice of either gender (aged 8-10 weeks) were used as recipients of yellow-green microsphere (polystyrene FluoSpheres™, 505ex/515em nm, Invitrogen, USA) transplantation. Postnatal day 3-6 (P3-6) mice expressing fused human rhodopsin-GFP (*Rho*-GFP^*+*^ mice, kind gift of Dr. T. Wensel, Baylor College of Medicine, TX) were used as photoreceptor cell donors. Adult (6-8 weeks of age) immune-deficient *NOD.CB17-Prkdc^scid^/J (NOD/SCID)* mice and retinal-degenerate *C3H/HeJ-Pde6b^rd1^ (rd1)* mice of either gender were used as recipients of *Rho*-GFP^*+*^ photoreceptor cell grafts. All recipients were obtained from Jackson Laboratory (USA). All mice were housed in cages under a 12:12-hour light-dark cycle with water and food provided *ad libitum*.

### Donor cell and sheet collection

Donor photoreceptor cells or sheets were obtained from *Rho*-GFP^*+*^ mice (P3-6) as reported previously ^1^. Briefly, the cornea was cut along the limbus and the lens and vitreous body were removed. The neural retina was gently isolated and washed in sterile phosphate-buffered saline (PBS, Gibco, USA). To obtain donor cell suspensions, neural retinas were digested at 37°C for 9 minutes in papain solution and single cells were obtained following manufacturer instructions (Papain Dissociation System, Worthington Biochemical, USA). Living cells were counted using a hemocytometer after trypan blue staining and resuspended in PBS at a density of 1×10^5^ cells/μl. To obtain donor retinal sheets, primary neural retinas were cut into 1 x 1 mm^2^ or 1 x 2 mm^2^ sheets using 27-gauge (G) horizontal curved scissors (VitreQ, USA) under a dissection microscope. All grafts were transplanted within 3 hours of isolation.

### Transplantation

Yellow-green microspheres were transplanted into single or multiple sites including the vitreous cavity (VC), subretinal space (SRS), and intra-retinal (InR) in wild-type *C57/BL6J* mice (n=3 per group). Surgical procedures were performed as previously reported ^1,23^. Briefly, recipient mice were anesthetized with intraperitoneal injection of ketamine (100 mg/kg body weight) and xylazine hydrochloride (20 mg/kg body weight). Pupils were dilated with 1% (wt/vol) tropicamide (Bausch & Lomb, USA) to facilitate transpupillary visualization under an operating microscope (Leica, USA). For SRS transplantation, 2μl of microsphere suspension (in PBS) was tangentially injected into the SRS through the sclera using a 34G microneedle and syringe (Hamilton, USA) under direct vision. After the entire bevel was inserted into the SRS, a consistently sized retinal bleb was observed post injection. For VC transplantation, 3 μl of microsphere suspended in PBS was injected into the VC approximately 1 mm posterior to the corneal limbus using the 34G microneedle and Hamilton syringe. For multi-site transplantation, microspheres were transplanted into the VC, SRS, and InR locations using the same 34G microneedle. *Rho*-GFP^*+*^ cell suspension was transplanted into the SRS of either *NOD/SCID* mice (n=5) or *rd1* mice (n=9) following the protocol mentioned above. For retinal sheet transplantation, each sheet was placed into the bevel of a 26G microneedle with the photoreceptor side facing down and gently aspirated into the attached syringe, then injected into the SRS of *NOD/SCID* mice (n=5) following the protocol as described. In each case, successful injection was verified by direct visualization through the dilated pupil of the recipient.

### Multimodal cSLO imaging

Recipients with severe cataract (> 5% dilated pupillary cover) were excluded. Multimodal cSLO imaging was performed on wild-type recipients at intervals up to 3 months (n=9), on *NOD/SCID* recipients up to 1 week (n=2), up to 1 month (n=2) or 2~3 months (n=6), and on *rd1* recipients up to 2 months (n=9) post-transplantation. Recipients were anesthetized by intraperitoneal injection of ketamine (100 mg/kg body weight) and xylazine hydrochloride (20mg/kg body weight). Pupils were dilated with 1% (wt/vol) tropicamide eye drops. A mouse contact lens (PLANO, UK) was placed on the cornea to prevent corneal dehydration. Standard 55° multimodal imaging was performed using the cSLO system (Heidelberg Engineering, USA). SWF imaging (488/500nm excitation/emission with automatic real-time (ART) averaging of 52 frames) was used to visualize fluorescence from yellow-green microspheres or GFP^+^ cells. MR images focusing on the mid-VC, retina blood vessel layer (RBVL), and SRS planes were obtained at the approximate same area as the SWF signal. A color-balanced MR image was acquired in each case by three lasers, namely, blue reflectance (BR) (λ = 488 nm), green reflectance (GR) (λ = 515 nm), and infrared (IR) (λ = 820 nm) reflectance (>20 frames of ART averaging). SD-OCT line scans through the same area were recorded at a mean of 9 frames per B-scan using ART mode. Non-injected eyes served as blank controls for all imaging modalities.

### Quantification of multimodal cSLO imaging biomarkers

Multimodal cSLO imaging biomarkers were quantified using ImageJ ^24^. The fluorescent signal *size* and *intensity* of each graft were quantified. (1) The SWF image was converted to greyscale (8 bit) and the scale bar was deleted to avoid its interference with overall intensity estimation; (2) The detection threshold was set to detect the entire graft and the percentage of selected pixels was recorded; (3) The fluorescent signal size and intensity of each graft were calculated as follows: To determine size, the numerical ratio of the graft area to total imaged fundus area was calculated. To adjust for variations in image exposure, the ratio of graft intensity to background intensity was calculated. For *graft length* quantification, the horizontal SD-OCT line scan through the maximum graft dimension was selected manually. The maximum graft length was measured using the caliper tool (dimension A). The recipient retinal thickness was measured as the distance from the RNFL to the retinal pigment epithelium (RPE) band (dimension B). Dimensions A and B were converted from inches to μm. Relative graft length was calculated from the ratio of graft length (μm) to recipient retinal thickness (μm) to standardize the scoring by controlling for the variability in retinal thickness at each site. The sizes of opacities presumed to be retinal *hemorrhages* were quantified by determining the ratio of hemorrhage area to total imaged fundus area on the MR image. The hemorrhage area was delineated by freehand selection. The total imaged fundus area was selected by threshold adjustment. To quantify the degree of recipient *retinal atrophy*, the thickness of the thinnest retina above the graft and a non-grafted control region were measured. The ratio of above-graft retina thickness to control retina was calculated as a measure of retinal atrophy. *Graft placement, lamination*, and *pre-retinal proliferation* biomarkers were manually determined based on SD-OCT image data.

### Histology

After sacrifice, perfusion fixation was performed with 4% paraformaldehyde (PFA) (Electron Microscopy Sciences, USA) in PBS. Eyes were removed and placed in 4% PFA/PBS for one hour and dehydrated in a sucrose gradient (10%, 20%, 30%), then blocked in optimal cutting temperature compound (Sakura Finetek, USA). Cryosections (10 μm) were cut by microtome (CM 1850, Leica, USA) and affixed to Superfrost Plus microscope slides (Fisher Scientific, USA) for staining. Sections of transplanted *rd1* mice eyes (n=6) were rinsed with PBS (5 min x 2), then permeabilized and blocked using a mixture of 0.1% Triton-X100 and 5% goat serum in PBS for 1 hour at room temperature (RT). Sections were rinsed with PBS (5 min x 3) and incubated with primary antibody overnight at 4°C. Primary antibodies used were goat anti-GFP (FITC) (1:200), rabbit anti-REC (1:1000), and rabbit anti-Pde6β (1:400), all from Abcam (USA). After rinsing in PBS (5 min x 3), sections with rabbit primary antibody were incubated with 1:500 goat-anti-rabbit Cy3-tagged secondary antibody (Abcam, USA) for one hour at RT, rinsed in PBS (5 min x 3) and counterstained nuclear with 4’, 6-diamidino-2-phenylindole (DAPI, Sigma, USA), rinsed in PBS (5 min x 2) and mounted using ProLong Diamond mounting media (Life Technology, USA). Seven equally distributed sections per eye were imaged under a confocal laser scanning microscope (LSM 510, Zeiss, USA). Positively stained cells were manually counted using ImageJ.

### Statistical analysis

Grading and IHC staining data were analyzed using the nonparametric Kendall’s tau-b correlation (two-tailed). Statistical analysis was carried out using SPSS software (version 25.0; SPSS, Inc., Chicago, IL, USA). *P*<0.05 taken as significant. Correlation coefficient (τ_b_) > 0.7 was considered as high correlation. Correlation graphs were drawn with GraphPad Prism software (version 7, CA, USA).

## Results

### Location detection using dynamic imaging with depth of focus modulation

Anatomical targets for cell delivery in animal models include the subretinal space (SRS) and the vitreous cavity (VC). *In vivo*, the SRS and VC are separated in depth. We aimed to evaluate the utility of *in vivo* multimodal cSLO imaging in differentially detecting the depth location of transplanted cells at these sites – and therefore, if multimodal cSLO imaging could be used to detect off-target delivery and/or cell migration, from the SRS to the VC or vice-versa, over time. We also aimed to understand whether depth detection accuracy of individual cells would be affected if the cells were present at different depth locations simultaneously. For example, cells could be found at different depth locations after complicated or traumatic delivery, wherein cells could be deposited in the VC (off-target) in addition to on-target SRS delivery. We established a noncellular model for initial imaging studies by using fluorescent yellow-green microspheres, instead of living fluorescence-labeled cells, as the delivery substrate. This model enabled us to address the technical questions above while avoiding challenges related to donor cell supply, immune cell rejection, and donor cell death. Fluorescent yellow-green microspheres were delivered into either the SRS, or the VC, in separate wild-type eyes as depth location controls. In test eyes, fluorescent yellow-green microspheres were delivered at multiple depth locations in the same eye (e.g. SRS, VC, and InR simultaneously), by gradually expelling the microspheres while the delivery needle was being translated axially.

Imaging with SWF, MR, and OCT modes was conducted one week after transplantation, giving sufficient time for the retinal bleb from SRS delivery to resolve, and the SRS cells to settle into a planar distribution. Dynamic manual focal plane modulation was performed to capture images at different focal depths in the eye, by adjustment of the *Z*-position using the *Z* micromanipulator. Images were acquired at the focal plane of the retinal blood vessel layer (*Z*_RBVL_) and at the focal plane of the subretinal space (*Z*_SRS_). Confirmation of the focal plane was obtained by noting that the retinal blood vessels were in sharp focus at *Z*_RBVL_, but were indistinct at *Z*_SRS_.

With SWF imaging, intraocular microspheres were detectable as bright objects scattered across the field of view. Depth-specific information of the transplanted microspheres could not be reliably obtained on static or dynamic (with focal plane modulation) imaging in the SRS control, VC control, or test case (Fig.1 A1, B1, C1).

**Figure 1.**
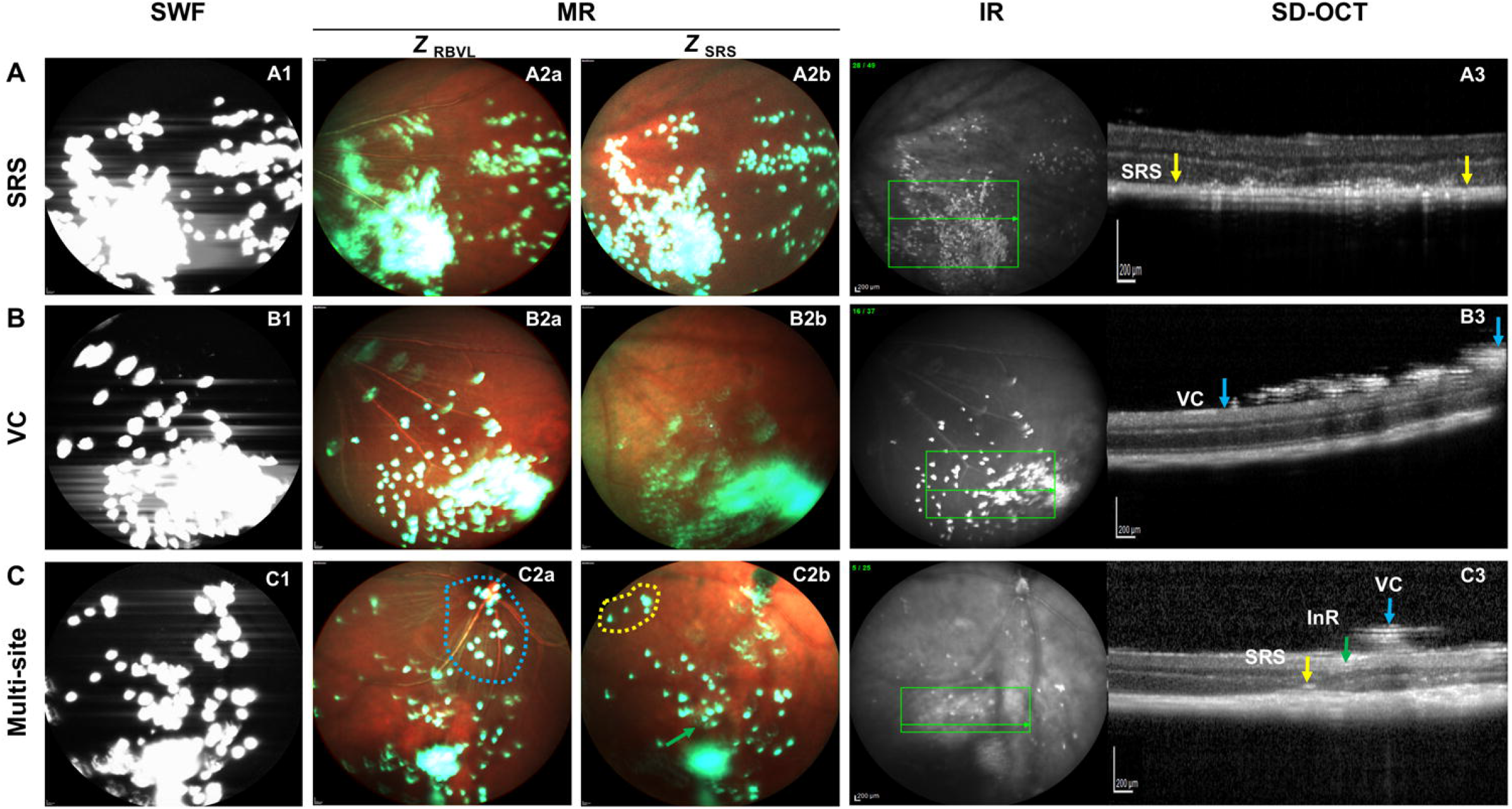
Location detection using dynamic imaging with depth of focus modulation in wild-type mice at 1-week post-transplantation. (A) Imaging of SRS-transplanted microspheres using SWF (A1), MR (A2a-2b), IR and SD-OCT (A3). (B) Imaging of VC-transplanted microspheres using SWF (B1), MR (B2a-2b), IR and SD-OCT (B3). (C) SWF (C1), MR (C2a-2b), IR and SD-OCT (C3) imaging of a test eye transplanted microspheres at multiple depth locations, including SRS, VC, and InR. All MR imaging were modulated dynamically at the focal plane of the retinal blood vessel layer (Z_RBVL_) or at the focal plane of the subretinal space (Z_SRS_). Yellow dashed circle and yellow arrow: microspheres located in SRS; Blue dashed circle and blue arrow: microspheres located in VC; Green arrow: microspheres located in InR. *Abbreviations: SRS: sub-retinal space; VC: vitreous cavity; InR: intra-retina; RBVL: retina blood vessel layer; SWF: short-wavelength fluorescence; MR: multicolor reflectance; IR: infrared reflectance; SD-OCT: spectral domain optical coherence tomography*.

With MR imaging, depth location of transplanted microspheres could easily be detected using dynamic imaging in the controls, where all microspheres were at one of two possible depths in each eye (SRS or VC). In the SRS location control, SRS microspheres were out of focus at *Z*_RBVL_ (Fig.1 A2a) and were in better focus at *Z*_SRS_ (Fig.1 A2b). In the VC location control, VC microspheres were in better focus when the focal plane set at *Z*_RBVL_ (Fig. 1 B2a) rather than at *Z*_SRS_ (Fig.1 B2b).

In test eyes with intentional multi-site delivery, the depth-specific information of some, but not all, microspheres could be obtained using dynamic focal plane modulation. Microspheres that were in sharp focus at *Z*_RBVL_ plane were taken to be at the VC location (Fig.1 C2a), whereas those that were in sharp focus at *Z*_SRS_ were taken to be at the SRS location (Fig.1 C2b). Several microspheres (single or clumped) did not change in sharpness appreciably from focal plane modulation and appeared similarly indistinct at *Z*_RBVL_ and *Z*_SRS_. These were presumed to be located at an intermediate depth location between the SRS and the VC, i.e., at the InR location (Fig.1 C).

SD-OCT imaging provided additional information regarding the microsphere depth location. SD-OCT imaging was a more direct assay of graft signal depth. However, only small areas could be sampled at one time and the data were generated as line scans. This made it challenging to integrate graft depth location information efficiently across the entire volumetric topography of the posterior eye, in comparison to MR imaging. Both SD-OCT and MR modes faced the limitation that axially superimposed microspheres were obscured, or in shadow, in the viewing plane. Nevertheless, SRS (Fig.1 A3) and VC (Fig.1 B3) microspheres were still separately distinguishable. InR microspheres were not clearly distinguishable from SRS and VC microspheres on SWF or MR imaging. The location of InR microspheres was identified more clearly using SD-OCT imaging (Fig.1 C3).

### Longitudinal multimodal cSLO detection of microsphere migration

Multimodal cSLO imaging detected the migration of both SRS and VC microspheres over three months of observation. SWF imaging showed that several clusters appeared to have dispersed over this time period, while other clusters appeared to have coalesced (Fig. 2 A1a–A1b and B1a–B1b). Whether these microspheres that changed location or distribution were in the SRS or VC was not clear from SWF data.

**Figure 2.**
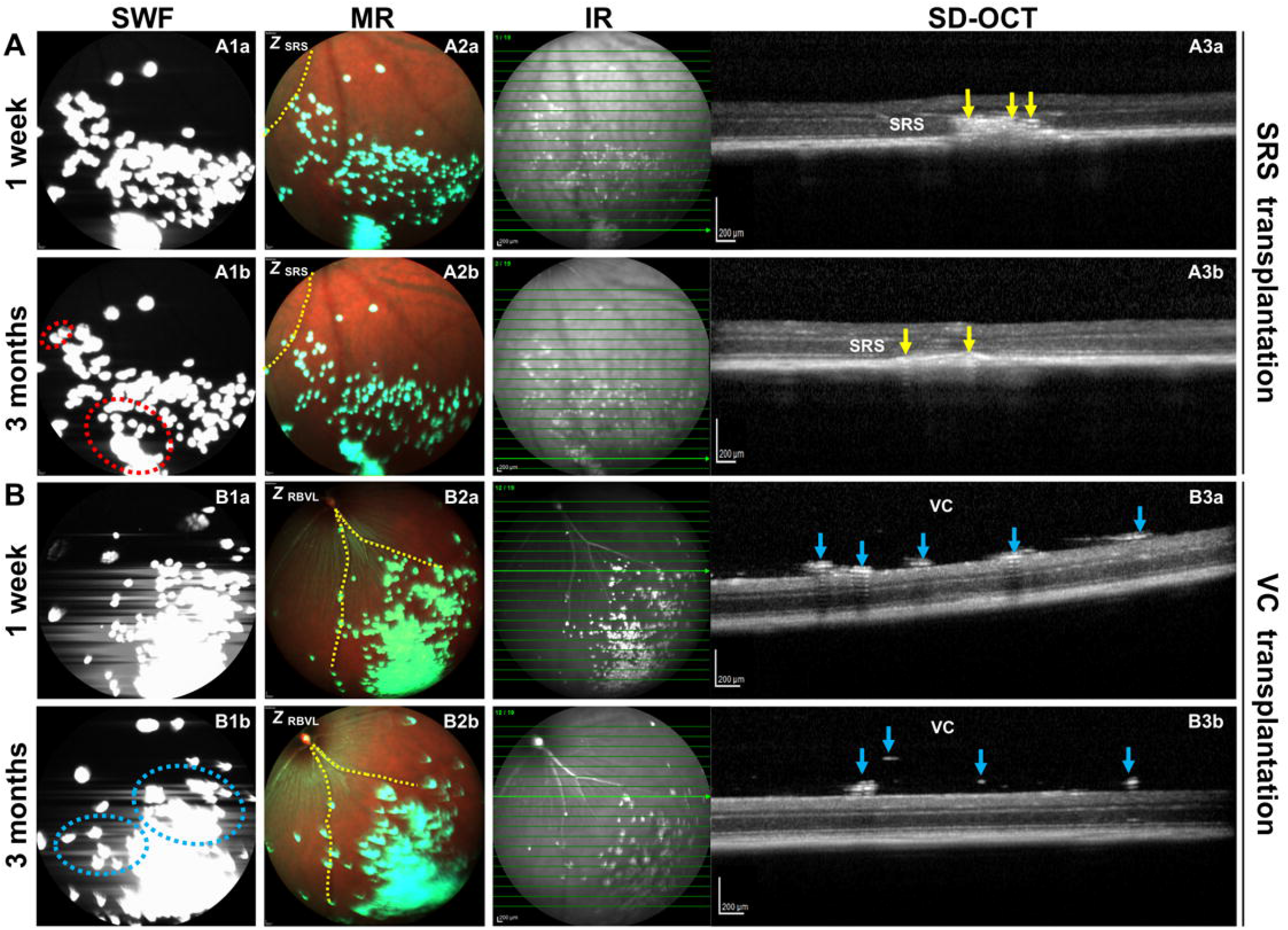
Longitudinal multimodal cSLO tracking the migration of transplanted microspheres in wild-type mice from 1 week to 3 months. (A) Multimodal cSLO imaging-based tracking of subretinally-transplanted microspheres. The SWF images (A1a-1b) shows the distribution changes of the microsphere clusters (red dashed circles). Migration of the microspheres was present on MR images (A2a-2b) with reference to the pattern of the retinal blood vessels (dashed yellow line). SD-OCT (A3a-3b) detected the migration of the microspheres in SRS (yellow arrows). (B) Multimodal cSLO imaging-based tracking of intravitreally-transplanted microspheres. The SWF images (B1a-1b) presented the distribution change of the microsphere clusters (blue dashed circles). Migration of the microspheres was detected on MR imaging (B2a-2b) with reference to the retinal blood vessels (yellow dashed line). SD-OCT (B3a-3b) showed the migration of VC microspheres from 1 week to 3 months (blue arrows). *Abbreviations: SRS: sub-retinal space; RBVL: retinal blood vessel layer; VC: vitreous cavity; SWF: short-wavelength fluorescence; MR: multicolor reflectance; IR: infrared reflectance; SD-OCT: spectral domain optical coherence tomography*.

The migration of SRS microspheres over three months of observation, even across relatively small distances, could be discerned using MR imaging with reference to the pattern of retinal blood vessels (Fig. 2 A2a–A2b). The majority of VC microspheres that migrated appeared to do so in an inferior direction, likely under the influence of gravity (Fig. 2 B2a–B2b). SD-OCT also showed SRS and VC microsphere migration over this time period, although exact co-registration of imaging across time points was challenging (Fig. 2 A3a–A3b and B3a–B3b).

### Longitudinal multimodal cSLO tracking of *Rho*-GFP^+^ photoreceptor cells

Based on the positive results in the non-cellular model above, we proceeded to investigate the utility of multimodal cSLO imaging in tracking longitudinally transplanted *Rho*-GFP^+^ photoreceptor cells. Multimodal cSLO imaging of recipient eyes was performed serially for up to three months after transplantation. *Rho*-GFP^+^ grafts (including cell suspension and sheet grafts) were detected in 13/19 eyes during short-term (1–4 weeks), and in 8/15 eyes during long-term (2~3 months), observation. Representative cSLO images from a *rd1* recipient, transplanted with *Rho*-GFP^+^ photoreceptor cell suspensions, are shown in Fig.3. By SWF imaging, the majority of transplanted cells appeared to be organized in clumps, while several others were more dispersed, which indicated scattered cells (Fig.3 A1). MR imaging (focused at *Z*_SRS_) was less effective at detecting cells in the same transplanted area (Fig.3 A2). The BR, GR, and IR channel signal patterns partially overlapped with each other (Fig.3 A2). Areas that were detectable on each MR imaging channel assumed a pattern that did not match the SWF pattern (Fig.3 A2). This potentially indicated differing anatomical correlates for each imaging modality signal. SD-OCT scans through the area of bright SWF signal presented reflective SRS cells. Interestingly, MR imaging (composite image) showed a dark-red circle adjacent to the graft (Fig.3 A2, white arrow), that correlated with retinal thinning that was detected on SD-OCT imaging (Fig.3 A4). Significant changes in multimodal imaging signals were observed in the same retina at two months. The GFP signal significantly decreased in size, and intensity, on SWF image (as shown in Fig.3 B1). The MR-detectable area changed in comparison to the one-month MR image. Interestingly, SD-OCT imaging showed a slight reduction of SRS graft length, despite the significant reduction of GFP+ area and intensity (Fig. 3 B3 vs. A3). SD-OCT imaging also detected progression of retinal thinning (Fig.3 B4 vs. A4).

**Figure 3.**
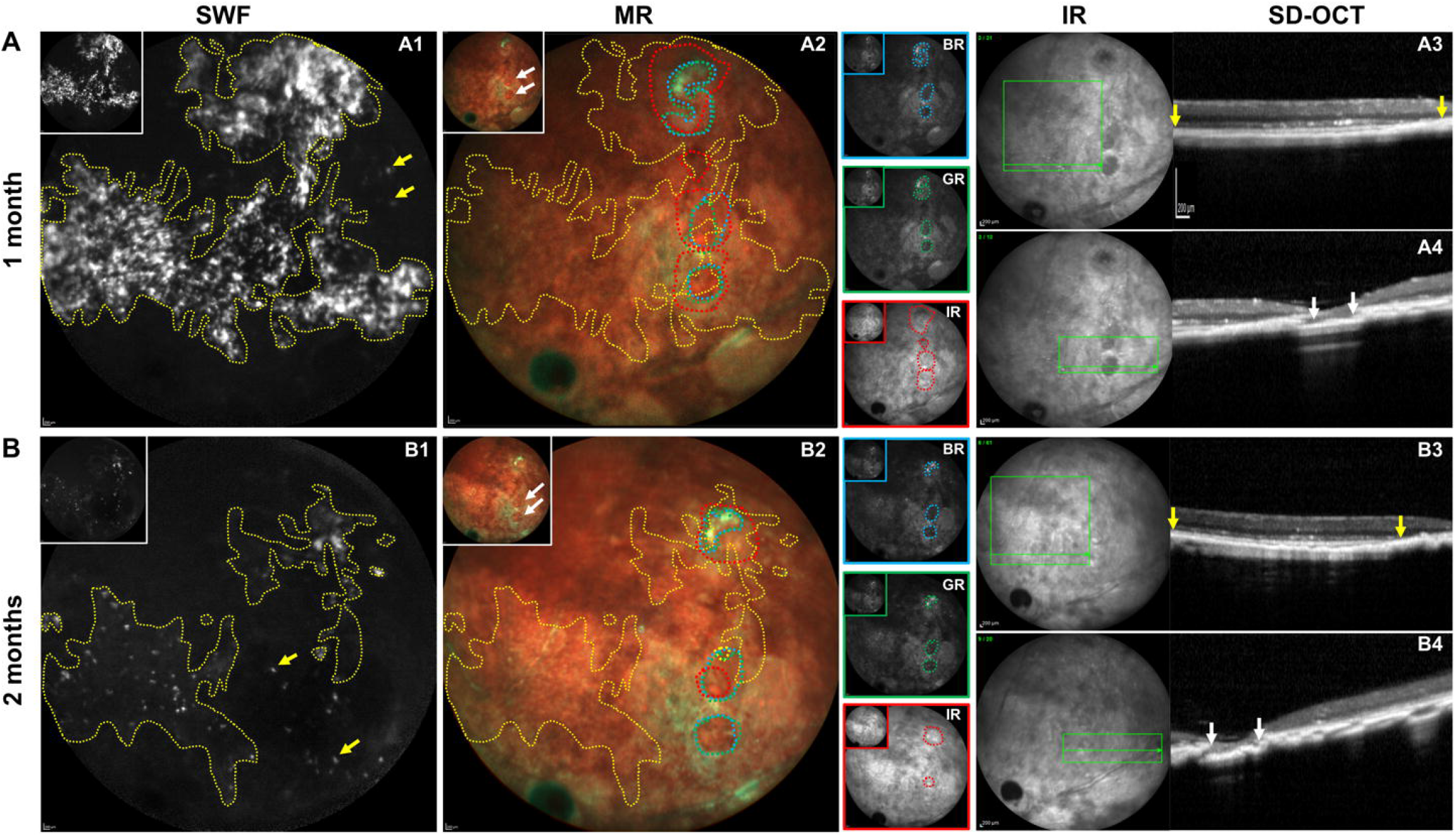
Longitudinal multimodal cSLO tracking of *Rho*-GFP^*+*^ photoreceptor cells *in vivo*. (A) Representative SWF (A1), MR (A2), IR fundus and SD-OCT images (A3-4) of a *rd1* recipient transplanted with *Rho*-GFP^*+*^ photoreceptor cell suspensions at 1 month post-transplantation. (B) Representative SWF (B1), MR (B2), IR fundus and SD-OCT images (B3–4) showing changes of the eye at 2 months post-transplantation. Original images without labeling are shown on the upper left corner of each annotated image. Yellow dashed circle: SWF-highlighted area; Blue-/green-/red-dashed circle indicate highlighted areas on BR, GR, IR images, respectively. Between yellow arrow: grafted cells; Between white arrow: recipient retinal atrophy. *Abbreviations: SWF: short-wavelength fluorescence; MR: multicolor reflectance; BR: blue reflectance; GR: green reflectance; IR: Infrared reflectance; SD-OCT: spectral domain optical coherence tomography*.

### Scoring of multimodal cSLO imaging biomarkers

A scoring scheme (Table 1) was developed to enable the description of changes quantitatively over time. The scoring scheme included the following indices: fluorescence signal *size* and *intensity* (SWF data); *graft length, graft placement*, and *lamination* (SD-OCT data); and complications, including *hemorrhage, recipient retinal atrophy*, and *peri-retinal proliferation* (MR and SD-OCT data).

**Table 1.**
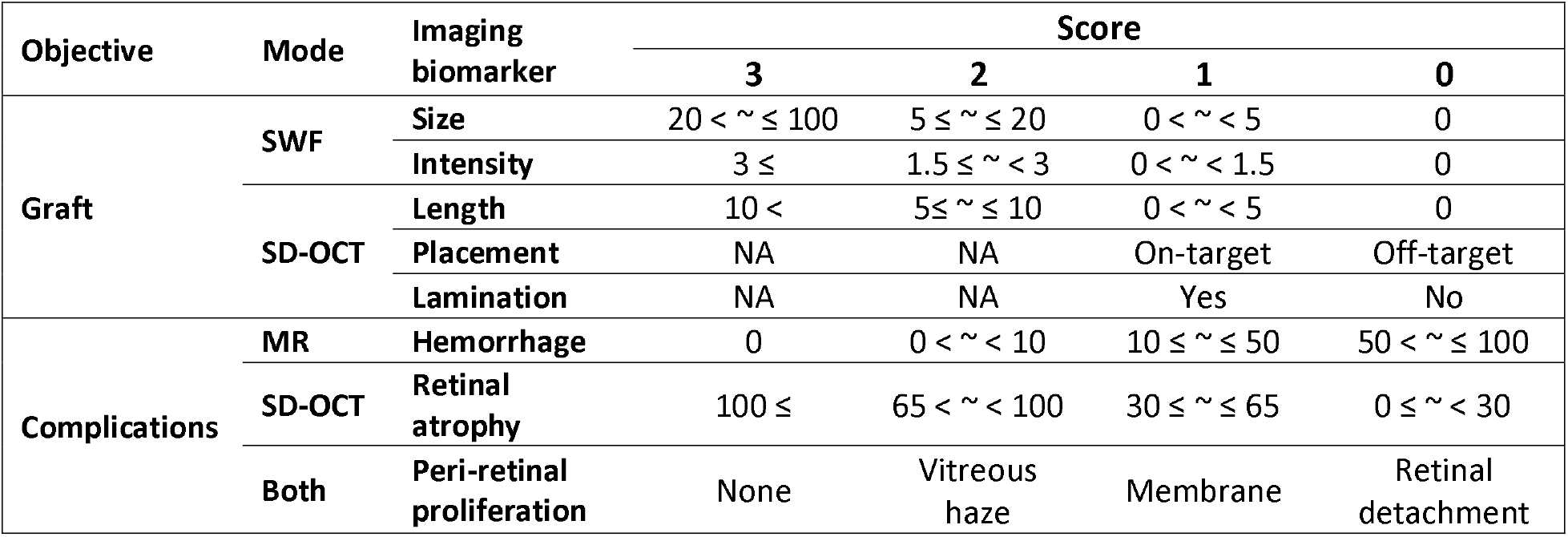
Scoring of multimodal cSLO imaging biomarkers. Imaging biomarkers were developed from multimodal imaging modes to describe graft status and recipient complications. For grafts, SWF imaging data indicated fluorescent *size* and *intensity*. SD-OCT imaging presented *graft length, graft placement* and intra-graft *lamination*. For complications, MR imaging present the *hemorrhage*. SD-OCT imaging detected the *retinal atrophy*. The *peri-retinal proliferation* was manually determined based on MR and SD-OCT image (“both” above) data. The scores of 0 to 3 were applied for each imaging biomarker as appropriate. *Abbreviations: NA: Not applicable*.

Fluorescence signal *size* was scored as 3 if the ratio of graft size to fundus area was > 20%, as 2 if the ratio was ≥ 5% and ≤ 20%, as 1 if the ratio was < 5% and > 0%, and as 0 if the ratio was equal to 0. Fluorescence *intensity* was scored from 0 to 3, with reference to the relative signal intensity of the graft versus background, as defined in Table 1. Fig. 4A shows representative images of transplanted *Rho*-GFP^*+*^ photoreceptor cells scored from 0 to 3 for signal size (a higher score indicated larger grafts), and signal intensity (a higher score indicated brighter GFP fluorescence).

**Figure 4.**
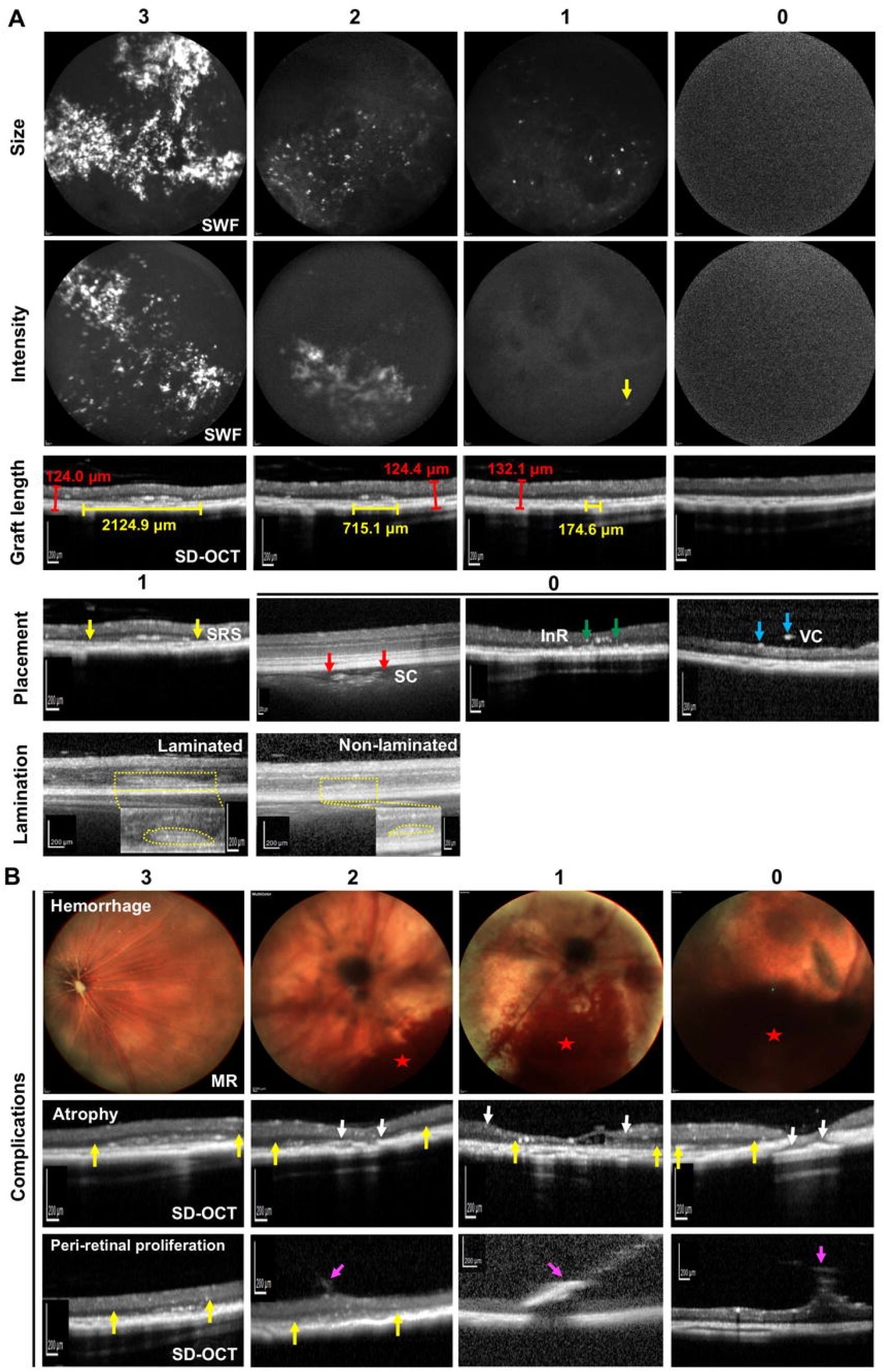
Representative images showing the scoring of multimodal cSLO imaging biomarkers. (A) Representative SWF and SD-OCT images of transplanted *Rho*-GFP^*+*^ photoreceptor cells. Image biomarkers include graft *size*, fluorescent *intensity, graft length, graft placement*, and intra-graft *lamination*. The scores were applied for each imaging biomarker as appropriate. (B) Representative MR and SD-OCT images of imaging biomarker complications, including *hemorrhage*, *recipient retinal atrophy*, and *peri-retinal proliferation*. A higher score indicated a less extensive or less severe complication. Between yellow arrows: grafts in sub-retinal space (SRS). Between red arrows: grafts in sub-choroid (SC). Between green arrows: grafts in intra-retina (InR). Blue arrows: grafts in vitreous cavity (VC). Between white arrows: recipient retinal atrophy. Magenta arrow: peri-retinal proliferation. Yellow dashed circle: laminated/non-laminated grafts. Yellow line: graft length. Red line: recipient retina thickness. Red star: hemorrhage. Note: the figure shown above for size score 3 is the same as Figure 3A.

*Graft length* score was assigned based on the maximum length of the graft on SD-OCT line scans. A score was assigned according to the ratio of graft length to the recipient retinal thickness reference within each image. A ratio of >10 was scored as 3, <5 was scored as 1, and a ratio between those values was scored as 2. Score 0 was assigned when no graft was found on SD-OCT images (Table 1). For example, the length of grafted photoreceptor cells in one transplanted eye was 2124.9 μm and the thickness of control recipient retina (in a proximal area without a graft) was 124.0 μm. The ratio of graft length to control recipient retinal thickness was 17.1. Therefore, it was assigned a graft length score of 3 (Fig. 4A). A higher graft length score indicated a longer maximum graft dimension in the horizontal axis.

The *graft placement* score was binary, based on SD-OCT imaging data. In the cell transplantation experiments, the intended graft location was the SRS. Therefore, a score of 1 was assigned if the graft was located entirely in the SRS, and 0 was assigned if any part of the graft was located outside the SRS, or if no graft was detected.

On the presumption that detectable intra-graft *lamination* was a favorable finding, which indicated the presence of layered cell arrangement in the graft, a lamination score of 1 was assigned to grafts in which the SD-OCT images showed a clear intra-graft lamination pattern (Fig.4A).

Complications – namely, *hemorrhage, retinal atrophy*, and *peri-retinal proliferation* – that could adversely affect transplantation outcomes were scored based on their extent and severity. We used a higher score to indicate a less extensive, or less severe, complication as listed in Table 1. Representative images are shown in Fig. 4B.

### Assessment of longitudinal changes in imaging scores

We aimed to determine if the scoring scheme could detect changes in graft and recipient status over time in photoreceptor cell transplants (n=16 eyes with available longitudinal data). A score change in at least one biomarker was detected in 14 of 16 eyes, over the period of observation after transplantation (Table 2). The *size* score decreased in 8/16 eyes (score change of −1 in four eyes and −2 in four eyes) and did not change in 8/16 eyes. The *intensity* score decreased in six eyes, remained stable in eight eyes and increased in two eyes. The *graft length* score changed in 5/16 eyes (score change of +1 in one eye, −2 in two eyes, −1 and −3 in one eye for each). The *graft placement* score remained stable (score = 1) in 15/16 eyes. Placement score dropped from 1 to 0 in one eye, in which no graft was visible during follow up imaging. The graft *lamination* score remained stable in 2/3 eyes transplanted with retinal sheets (score remained as 1 in two eyes and decreased from 1 to 0 in one eye). The *complication* score was stable in 9/16 eyes (score remained as 3 in two eyes, as 2 in four eyes, and as 1 in three eyes). Changes in complication score were detected in 7/16 eyes (score change of −2 in one eye, −1 in two eyes, and +1 in four eyes).

**Table 2.** Quantification of longitudinal changes in imaging scores of transplanted mice. Multifocal SLO imaging were performed longitudinally in sixteen *Rho*-GFP^*+*^ photoreceptor cell transplanted mice. Imaging biomarkers, including graft *size*, fluorescence *intensity, graft length, graft placement, lamination*, and *complication*, were scored in each mouse. Lamination scoring was applied to retinal sheet grafts (n=3 eyes) only, considering their initially laminated structures. The score changes of individual imaging biomarker were quantified from the observation start time (week 1 or month 1) to the termination (up to 3 months). The number of eyes showing each possible magnitude of score change is listed in the table. *Abbreviations: NA: Not applicable*.

| Score change<br>(n=16) | SWF |  | SD-OCT |  |  | MR & SD-OCT |
| --- | --- | --- | --- | --- | --- | --- |
|  | Size | Intensity | Length | Placement | Lamination | Complications |
| + 3 | 0 | 0 | 0 | NA | NA | 0 |
| + 2 | 0 | 0 | 0 | NA | NA | 0 |
| + 1 | 0 | 2 | 1 | 0 | 0 | 4 |
| 0 | 8 | 8 | 11 | 15 | 2 | 9 |
| - 1 | 4 | 2 | 1 | 1 | 1 | 2 |
| - 2 | 4 | 3 | 2 | NA | NA | 1 |
| - 3 | 0 | 1 | 1 | NA | NA | 0 |

### Integration of biomarker scores into a grading system

We combined the different biomarker scores into a single grade for each image, so that we could integrate multiple biomarker data points for each eye. We assigned a grade (I–IV, reflecting the best to the worst outcomes, Table 3) for each eye, based on the sum of individual imaging biomarker scores for that eye. For this purpose, the complication score for each eye was defined as the lowest score (corresponding to the most severe or extensive complication) of the three possible complication scores of hemorrhage, atrophy, and peri-retinal proliferation. We adopted this approach to numerically disadvantage the grafts with more severe, or more extensive, complications.

**Table 3.**
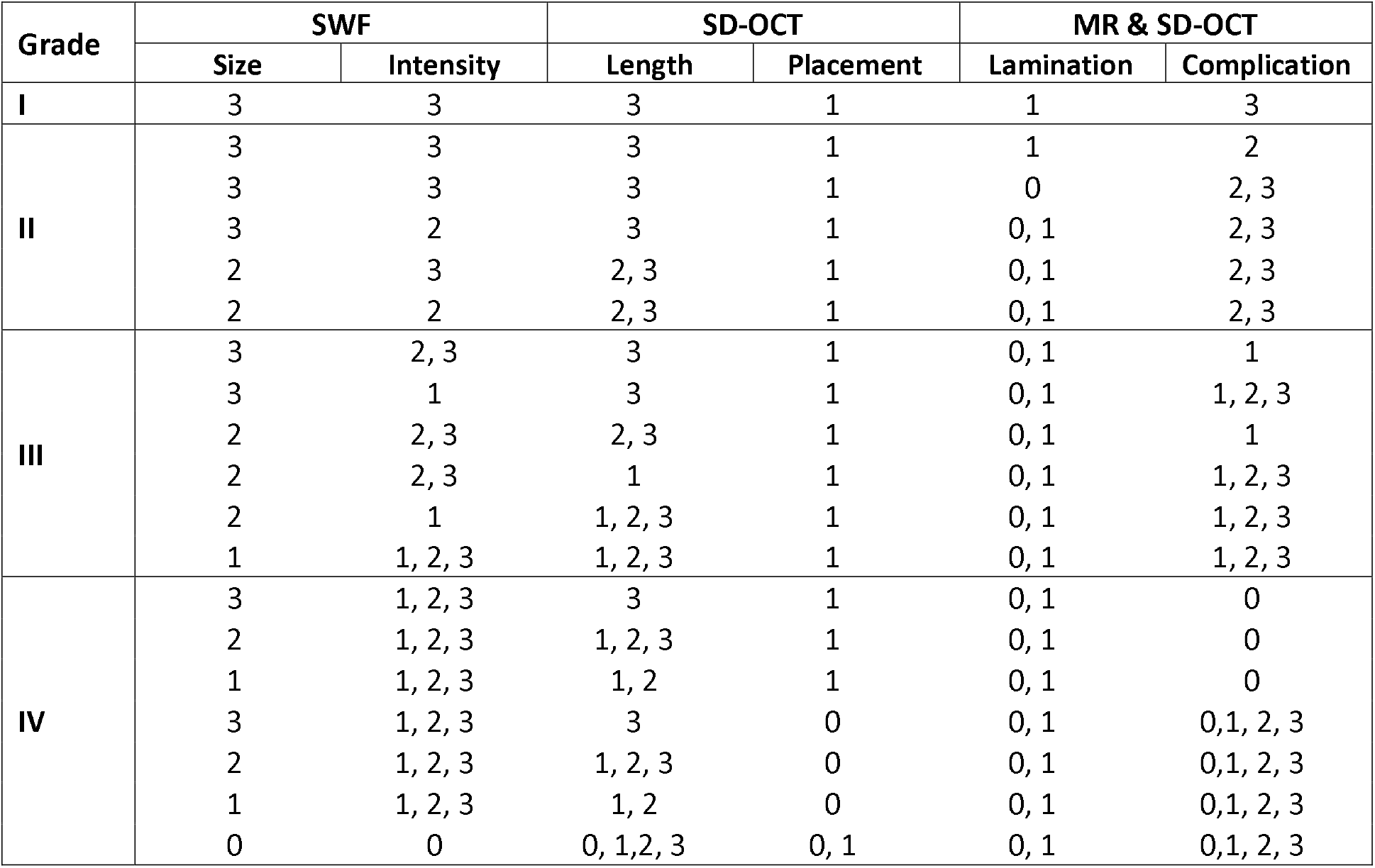
Integration of biomarker scores into a grading system. Grades I to IV were created by integrating the scores of individual imaging biomarkers, including graft *size*, fluorescent *intensity, graft length, graft placement, lamination* and *complication*. The lamination score was only applied to retinal sheet grafts. The combinations of the individual score are listed in the table.

| Grade | SWF |  | SD-OCT |  | MR & SD-OCT |  |
| --- | --- | --- | --- | --- | --- | --- |
|  | Size | Intensity | Length | Placement | Lamination | Complication |
| I | 3 | 3 | 3 | 1 | 1 | 3 |
| II | 3 | 3 | 3 | 1 | 1 | 2 |
|  | 3 | 3 | 3 | 1 | 0 | 2, 3 |
|  | 3 | 2 | 3 | 1 | 0, 1 | 2, 3 |
|  | 2 | 3 | 2, 3 | 1 | 0, 1 | 2, 3 |
|  | 2 | 2 | 2, 3 | 1 | 0, 1 | 2, 3 |
| III | 3 | 2, 3 | 3 | 1 | 0, 1 | 1 |
|  | 3 | 1 | 3 | 1 | 0, 1 | 1, 2, 3 |
|  | 2 | 2, 3 | 2, 3 | 1 | 0, 1 | 1 |
|  | 2 | 2, 3 | 1 | 1 | 0, 1 | 1, 2, 3 |
|  | 2 | 1 | 1, 2, 3 | 1 | 0, 1 | 1, 2, 3 |
|  | 1 | 1, 2, 3 | 1, 2, 3 | 1 | 0, 1 | 1, 2, 3 |
| IV | 3 | 1, 2, 3 | 3 | 1 | 0, 1 | 0 |
|  | 2 | 1, 2, 3 | 1, 2, 3 | 1 | 0, 1 | 0 |
|  | 1 | 1, 2, 3 | 1, 2 | 1 | 0, 1 | 0 |
|  | 3 | 1, 2, 3 | 3 | 0 | 0, 1 | 0, 1, 2, 3 |
|  | 2 | 1, 2, 3 | 1, 2, 3 | 0 | 0, 1 | 0, 1, 2, 3 |
|  | 1 | 1, 2, 3 | 1, 2 | 0 | 0, 1 | 0, 1, 2, 3 |
|  | 0 | 0 | 0, 1, 2, 3 | 0, 1 | 0, 1 | 0, 1, 2, 3 |

Grade I represented *highly favorable* transplantation outcomes, including a large graft (size score = 3, length score = 3), high GFP expression (intensity score = 3), SRS location (placement score = 1), presence of donor sheet lamination (score = 1), and no complications (score = 3). Grade II represented *favorable* transplantation outcomes with a score = 2 for any one of size, length, intensity, or complication, and placement score = 1, regardless of lamination score. Grade III represented *relatively unfavorable* transplantation outcomes with a score = 1 for any one of size, length, intensity, or complication, and placement score = 1, regardless of lamination score. Grade IV represented *unfavorable* transplantation outcomes with a score = 0 for any one of size, length, intensity, placement, or complication, regardless of lamination score.

### Application of the grading system to quantify longitudinal changes *in vivo*

To test the application of this system *in vivo*, we evaluated transplanted *Rho*-GFP^*+*^ photoreceptor cell suspensions and sheets over time. Follow-up data were available for 16 recipient eyes, that received either type of graft for up to three months post transplantation (one eye up to one month, nine eyes up to two months, and six eyes up to three months). Overall, a change in grade was found in 7/16 eyes. From week one to month one, 2/3 eyes remained at grade III, 1/3 eye dropped from grade III to grade IV. From month one to month two, 7/11 eyes remained at either grade III (2 eyes) or grade IV (5 eyes). A reduction in grade occurred in 4/11 eyes during this period, of which one eye dropped from grade I to II, two eyes dropped from grade II to III, and one eye dropped from grade II to IV. Among the six eyes that were tracked for three months, the grade was stable at II, III, and IV, from 1 month to 3 months in three eyes, respectively. Two eyes dropped in grade: one from II to IV and the other from III to IV. No eyes showed an increase in grade over time (Table 4).

**Table 4.**
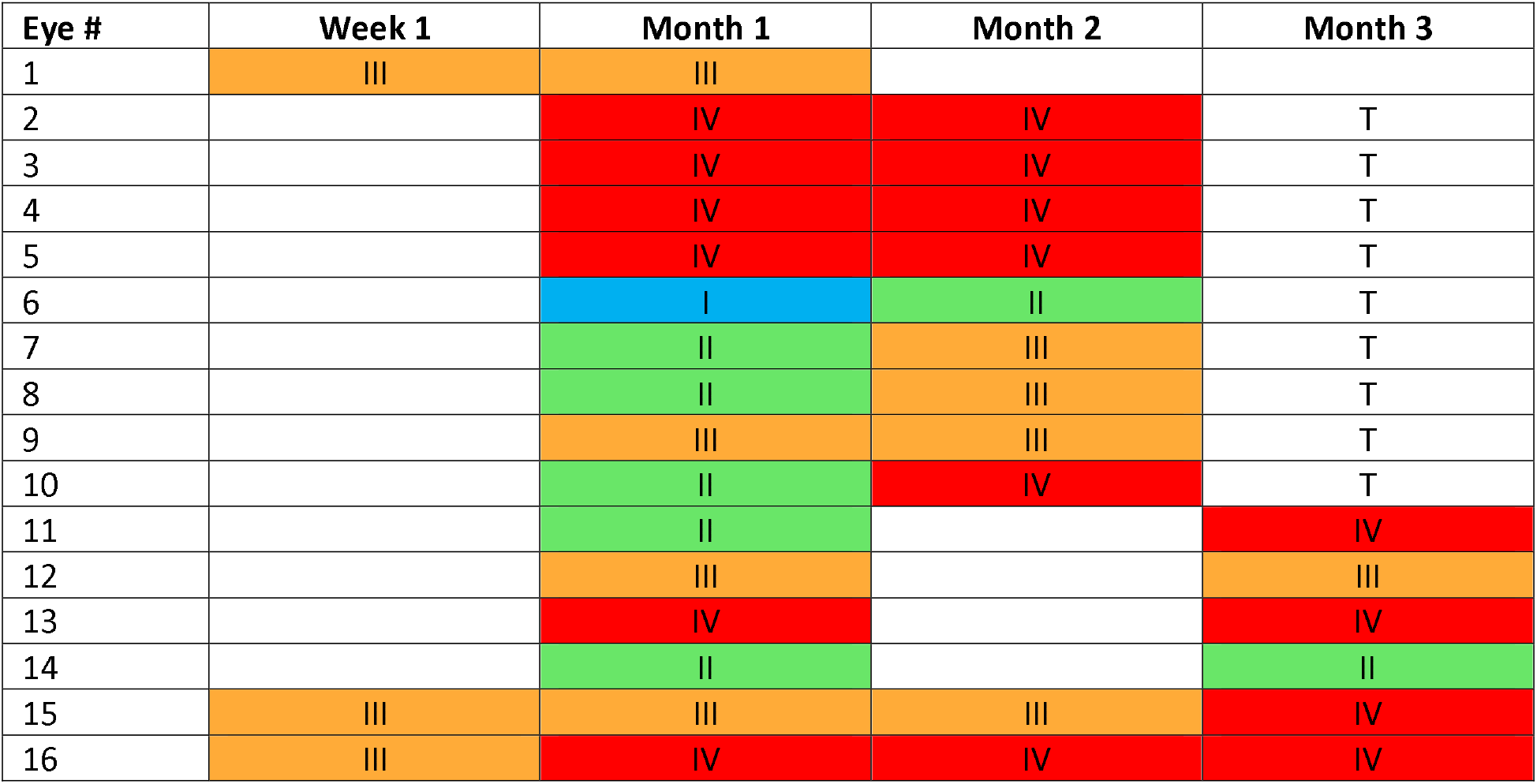
Application of the per-eye grading system in *Rho*-GFP^*+*^ photoreceptor cell transplanted mice. The grade of individual imaging biomarkers was quantified in 16 eyes for up to three months. Each grade was labeled as a specific color: Blue = Grade I; Green = Grade II; Orange = Grade III; Red = Grade IV. Grade change was found in 7/16 eyes in total. Blank cells indicate no data for that time point. *Abbreviation: T= assay terminated*.

| Eye # | Week 1 | Month 1 | Month 2 | Month 3 |
| --- | --- | --- | --- | --- |
| 1 | III | III |  |  |
| 2 |  | IV | IV | T |
| 3 |  | IV | IV | T |
| 4 |  | IV | IV | T |
| 5 |  | IV | IV | T |
| 6 |  | I | II | T |
| 7 |  | II | III | T |
| 8 |  | II | III | T |
| 9 |  | III | III | T |
| 10 |  | II | IV | T |
| 11 |  | II |  | IV |
| 12 |  | III |  | III |
| 13 |  | IV |  | IV |
| 14 |  | II |  | II |
| 15 | III | III | III | IV |
| 16 | III | IV | IV | IV |

An example of tracking *Rho*-GFP^*+*^ photoreceptor cell suspension graft in an *rd1* recipient is shown in Fig. 5A. SWF imaging showed a visible reduction of SWF signal, from month one to month two post-transplantation. Graft size and intensity were 32.7% and 5.4 respectively, at one month. Accordingly, each of these biomarkers received a score of 3. However, at two months, we observed reduced size (2.7%) and intensity (2.1), thus reducing the scores to 1 and 2 respectively. Graft length was scored as 3 at one month (graft length ratio 2832.1/135.8 = 20.9) and remained at score 3 at two months, despite the reduced density and intensity scores. Graft placement was scored as 1 at both time points, because the transplanted cells remained in the SRS. Complications were scored as 3, since none were detected at both time points. Taken together, the graft was assigned Grade I (highly favorable) at month one, but dropped to grade III (relatively unfavorable) at month two (Fig. 5A).

**Figure 5.**
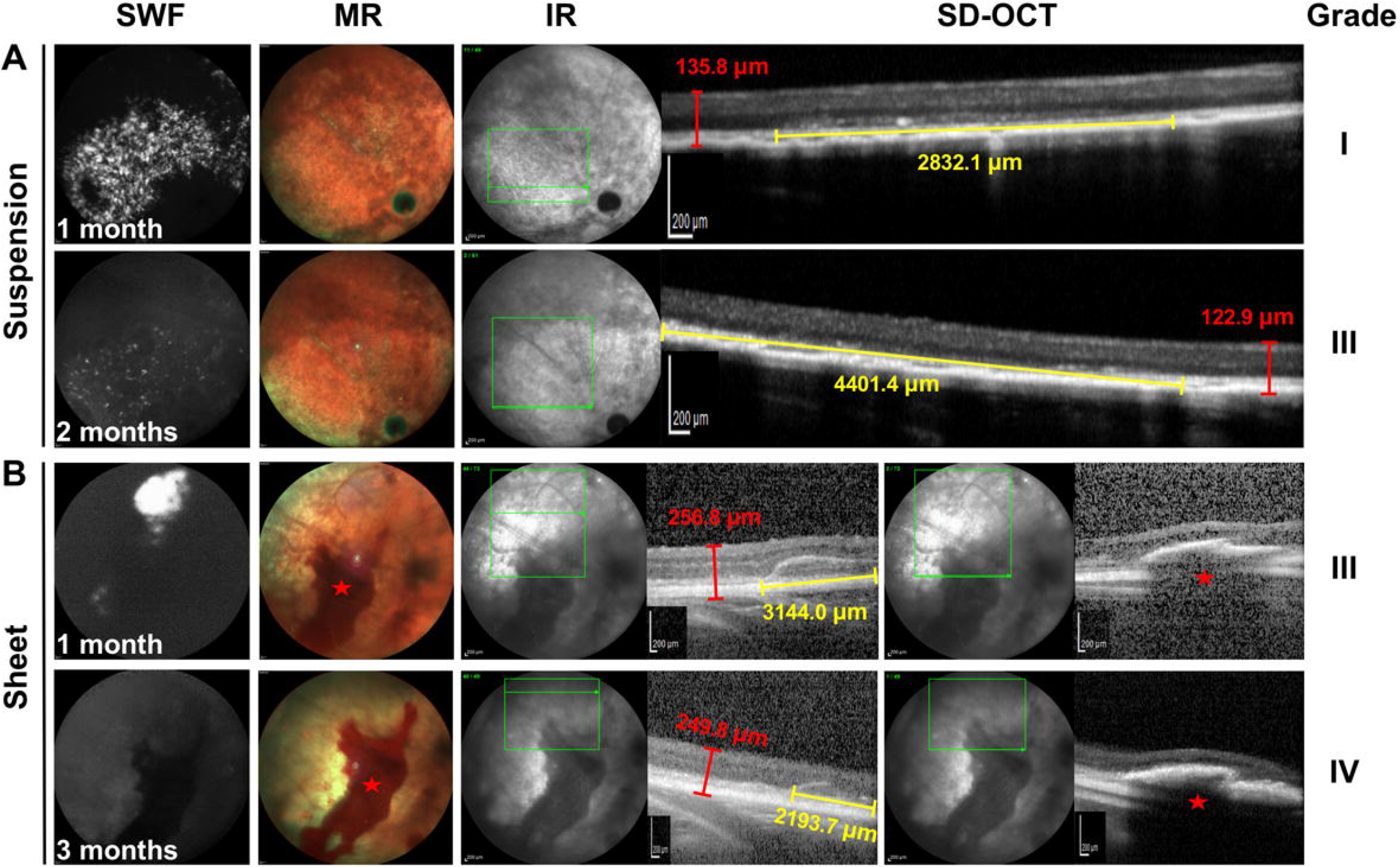
Application of the grading system to quantify longitudinal changes *in vivo*. (A) Multimodal cSLO images of a *rd1* mouse transplanted with *Rho*-GFP^*+*^ photoreceptor cell suspensions. The grade of imaging biomarkers changed from grade I to grade III over 2-month observation. (B) Multimodal cSLO images of a *NOD/SCID* mouse transplanted with *Rho*-GFP^*+*^ retinal sheets. The grade of imaging biomarkers changed from grade III to grade IV over 3-month observation. Yellow line: graft length; Red line: recipient retina thickness; Red star: hemorrhage.

An example of *Rho*-GFP^*+*^ retinal sheet transplantation in a *NOD/SCID* recipient is shown in Fig. 5B. At one month, GFP density (5.63%) was scored as 2, and intensity (5.27) was scored as 3. Graft length was scored as 3 (graft length ratio 3144.0/256.8 = 12.2). The graft remained located in the SRS (score 1) and showed an internal lamination pattern (score 1). For complications, a sizeable hemorrhage (22.4% area) was detected on MR imaging, and was thus scored as 1. Two additional complications (retinal atrophy and peri-retinal proliferation) both scored 3, because they were not detected. Therefore, the score of 1 from the hemorrhage, being the lowest score of the three potential complications, was selected as the final complication score. Three months later, no graft-related signal was detected on SWF image. Hence, graft size and intensity were both scored as 0. But the graft was still detectable on SD-OCT image, and the graft length score reduced to 2 (graft length ratio 2193.7/249.8 = 8.8). The graft was located in SRS, and so the placement was scored as 1. Graft lamination was scored as 0, because no intra-graft lamination was found on SD-OCT images. The complication score remained as 1, due to the persistent hemorrhage that had enlarged slightly (26.5% area). The grade of this graft changed from grade III (relatively unfavorable) to grade IV (unfavorable), over three months of tracking.

### Correlation of image grade with histological data

To test the extent to which the image-based grading system correlated with histological data, and its potential for use as a partial surrogate endpoint for histological analysis, we correlated the image grades and immunohistological findings in six *rd1* mice eyes transplanted with *Rho*-GFP^*+*^ photoreceptor cell suspensions.

The mouse presented in Fig. 5A was assigned grade III (relatively unfavorable outcome) at month two and was immediately sacrificed for IHC analysis. Numerous cells were located in the SRS, but few expressed Rho-GFP (Fig. 6A). These findings were consistent with the low GFP size (score = 1) and intensity (score = 2) scores, and the high graft length score (score = 3) (Fig. 5A). The low expression of photoreceptor markers, including Rho-GFP, recoverin (REC), and Pde6β, indicated relatively poor graft survival and maturation. Thus, the low grade of this mouse (grade III) correlated with the poor histological outcome. A similar correlation was found in five other tested eyes: more numerous GFP^+^, REC^+^, and Pde6β^+^ cells were found in specimens that were graded more favorably. Statistical analysis showed a high correlation between image grade and histology (τ_b_ >0.85, P<0.05, Fig. 6B).

**Figure 6.**
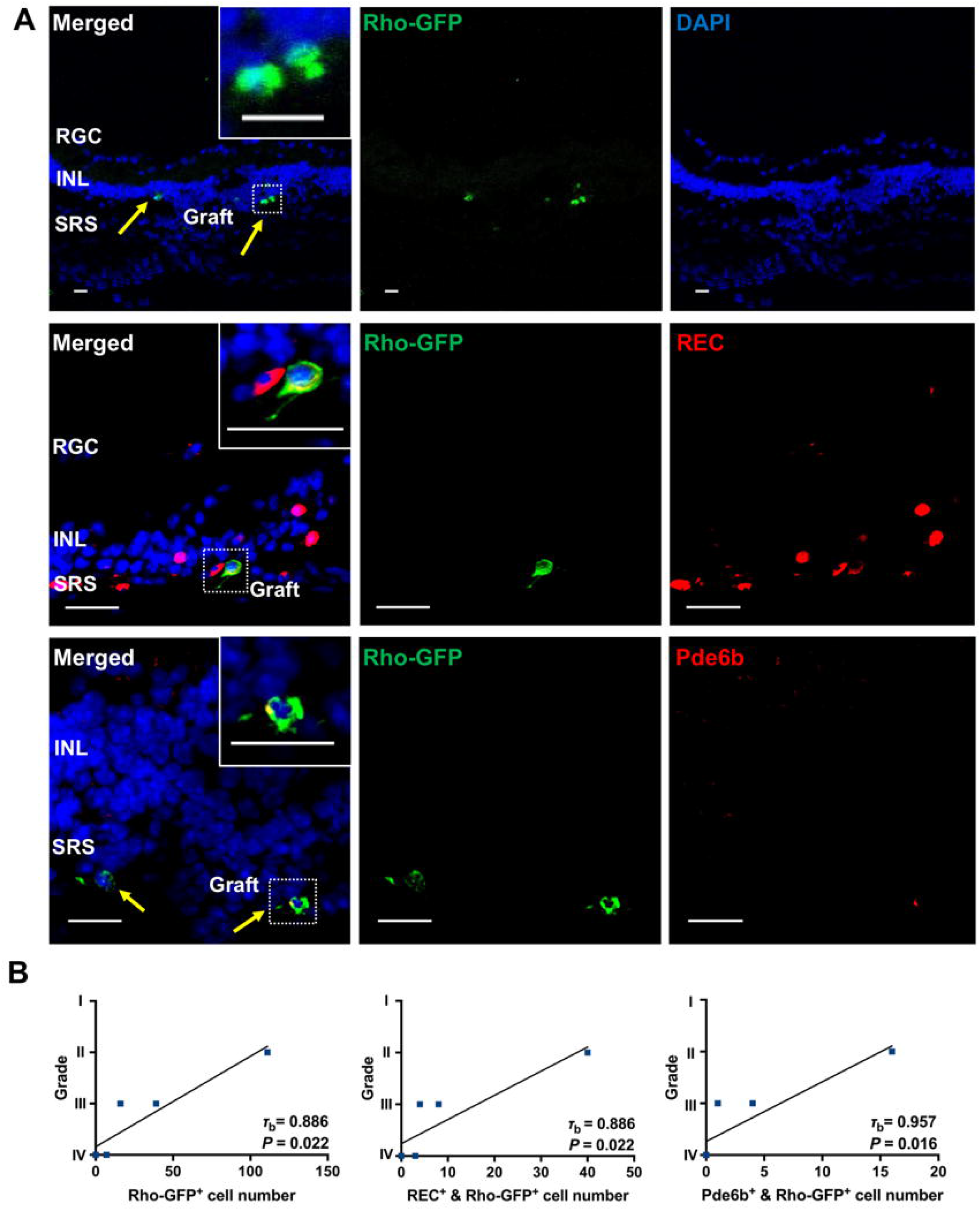
Correlation of imaging grade and histological data. (A) Representative immunohistochemistry (IHC) staining images of the *rd1* recipient in Fig. 5A. The mouse was transplanted with *Rho*-GFP^*+*^ photoreceptor cell suspension and was sacrificed 2 months post-transplantation for IHC staining. Photoreceptor-specific markers (Rho-GFP, REC, and Pde6β) were stained to evaluate graft survival and maturation. Magnified images are provided on the upper right corner. (B) Kendall’s tau-b correlation analysis of imaging grade and histological data in *rd1* mice (n= 6 eyes, 7 sections/eye): Rho-GFP positive-, REC and Rho-GFP double positive, and Ped6β and Rho-GFP double positive cells were counted and correlated with image grades. Yellow arrow: graft. τ_b_: correlation coefficient. Scale bar = 20 μm. *Abbreviations: RGC: retinal ganglion cell; INL: inner nuclear layer; SRS: sub-retinal space*.

## Discussion

The data show that multimodal imaging yielded a range of imaging biomarkers that were amenable to systematic quantitation. SWF detected graft size and GFP intensity, SD-OCT showed graft length, placement, and lamination, and MR reflectance and SD-OCT together showed complications including hemorrhage, atrophy, and peri-retinal proliferation. The imaging biomarkers could be scored and integrated into a per-eye grade, as a quantifiable and trackable measurement of transplantation outcomes over time. Confidence in the use of this system of tracking graft status was supported by its high correlation with histology.

Various cellular grafts (e.g. photoreceptor precursors, RPE cells, bone marrow cells, umbilical tissue cells, and others) have been studied in small and large animal transplantation models ^3,25–29^, *en route* to human application^30–34^. A typical goal in preclinical retinal cell therapy research is to assess the long-term structural outcomes (e.g. survival, maturation, and migration) of the transplanted cells in the context of functional outcome (e.g. visual function assays). Among the challenges is the fact that suboptimal grafts – i.e., those adversely affected by hemorrhage, off-target placement, reflux, retinal detachment, or retinal trauma – are only discovered by histological analysis at the end of the study period. This conventional process consumes time, manpower, and financial resources. Complicated grafts, while providing information on how and why transplants fail, do not typically contribute positively towards functional correlation.

Therefore, an *in vivo* imaging-based graft evaluation strategy could be useful to grade and stratify graft status, prior to targeted or stratified histological and functional analysis. Arguably, the most important structural aspect of a graft that should be ascertained prior to selection for functional correlation is whether on-target delivery was achieved completely. This information is not typically available in the living animal, prior to downstream terminal analyses, without imaging. There can be a high rate of off-target delivery in small animals, because of the challenge of precise maneuvers in small eyes, and also because nonvisual-guided delivery is commonly practiced. We found multimodal cSLO imaging to be very effective at determining graft placement in the SRS and VC, using a combination of information from SWF, MR, and SD-OCT. Interestingly, we detected the migration of transplanted material in the VC and SRS in several mice. This was likely due to animal movement, and the influence of gravity, where cells in the SRS, and more so in the fluid-filled VC, could shift over time while remaining in the same compartment.

cSLO fluorescence imaging has been used to detect red dye-labeled photoreceptor cells in the rat eye in short-term transplantation studies ^19,35^. However, fluorescence dye is not a reliable indicator of graft survival because it is non-specific to living cells and could also leak into recipient cells ^36^. We used human rhodopsin-fused GFP as the traceable donor-specific marker in this study. Using this system, graft survival and photoreceptor maturation could be indicated by GFP expression, because the GFP signals are restricted to relatively mature living Rho^+^ photoreceptor cells. In contrast to several prior studies, in which OCT imaging was used to ascertain graft survival ^20,37,38^, we used SWF imaging to track transplanted cells. We found that SWF imaging effectively presented the planar distribution of GFP+ grafts across the visible topographic extent of the recipient fundus, making it more efficient than OCT line scanning for graft detection ^39^.

It should be noted that MR imaging is based on reflectance, and not fluorescence. We found MR imaging to be relatively inefficient in detecting GFP+ cells compared to SWF imaging. Other possible explanations for this could lie in differences in the characteristics of microspheres and photoreceptors. Microspheres are larger imaging targets than photoreceptor cells (15 μm vs 1.2–1.4 μm, respectively). Fluorescence intensity in microspheres appears to be higher than that of GFP in photoreceptors, and also more uniform due to the lack of intercellular variation in expression level due to cell degeneration or other factors.

For a more complete interpretation of graft survival, we found it important to integrate information from SWF and SD-OCT modalities. Loss of SWF signal could be interpreted as graft degeneration. However, we often found SD-OCT data that were consistent with the continued presence of transplanted SRS material in those eyes. In most cases, histology showed that this material contained transplanted photoreceptor cells with downregulated GFP expression, likely accounting for their poor visibility on SWF imaging. Hence it is critical to consider the SWF graft size and intensity scores in the context of the SD-OCT graft length score, as photoreceptor grafts could manifest differentially in each modality.

Functional outcomes can also be influenced by complications that could degrade the graft. SWF, MR, and SD-OCT data were useful in detecting complications, such as hemorrhage, recipient atrophy, and peri-retinal proliferation. Changes in the number of RPE fluorophores ^40,41^, or the presence of absorbing/emitting material anterior to the RPE, will both produce fundus auto-fluorescence (FAF) signal changes on SWF images. Loss of signal could be caused by RPE atrophy ^42,43^, retinal edema ^44^, and hemorrhage ^45^, whereas increased signal could be associated with subretinal fluid ^46^ and drusen ^47,48^. Reduced FAF in areas of RPE atrophy has been reported in *rd1* mice ^42^. We found that visualization of the reduced FAF signal associated with atrophy was often obscured by the very bright GFP signal in transplanted eyes (Fig. 3A), so OCT imaging may be a more consistent method to detect RPE atrophy in such cases.

## Conclusions

Multimodal cSLO imaging biomarker analysis enabled quantifiable, and longitudinal, assessment of features of retinal regeneration by photoreceptor cell transplantation. Imaging biomarkers reflected graft survival and distribution, and recipient retinal complications. Multiple imaging biomarkers could be integrated into a per-eye grading scheme that enabled comprehensive tracking of changes in graft and/or recipient status in single eyes over time. Multimodal *in vivo* imaging of individual recipients in cohorts of mice may facilitate functional correlations, by enabling stratification of eyes according to accuracy of on-target placement, graft survival, graft size, and graft damage from complications. The application of systematic multimodal *in vivo* imaging could be useful in increasing the efficiency of preclinical retinal cell transplantation studies in rodents and other animal models, by reducing the reliance on end-point histology.

## Acknowledgments

We thank Dr. T. Wensel and Dr. Donald Zack for the kind gift of the *Rho*-GFP^*+*^ mice. We thank Diana Johnson and Dr. Karl A. Hudspith for their assistance.

