## Supplemental Figure S1 for "Quantifiable *In Vivo* Imaging Biomarkers of Retinal Regeneration by Photoreceptor Cell Transplantation"

**Supplementary Material**

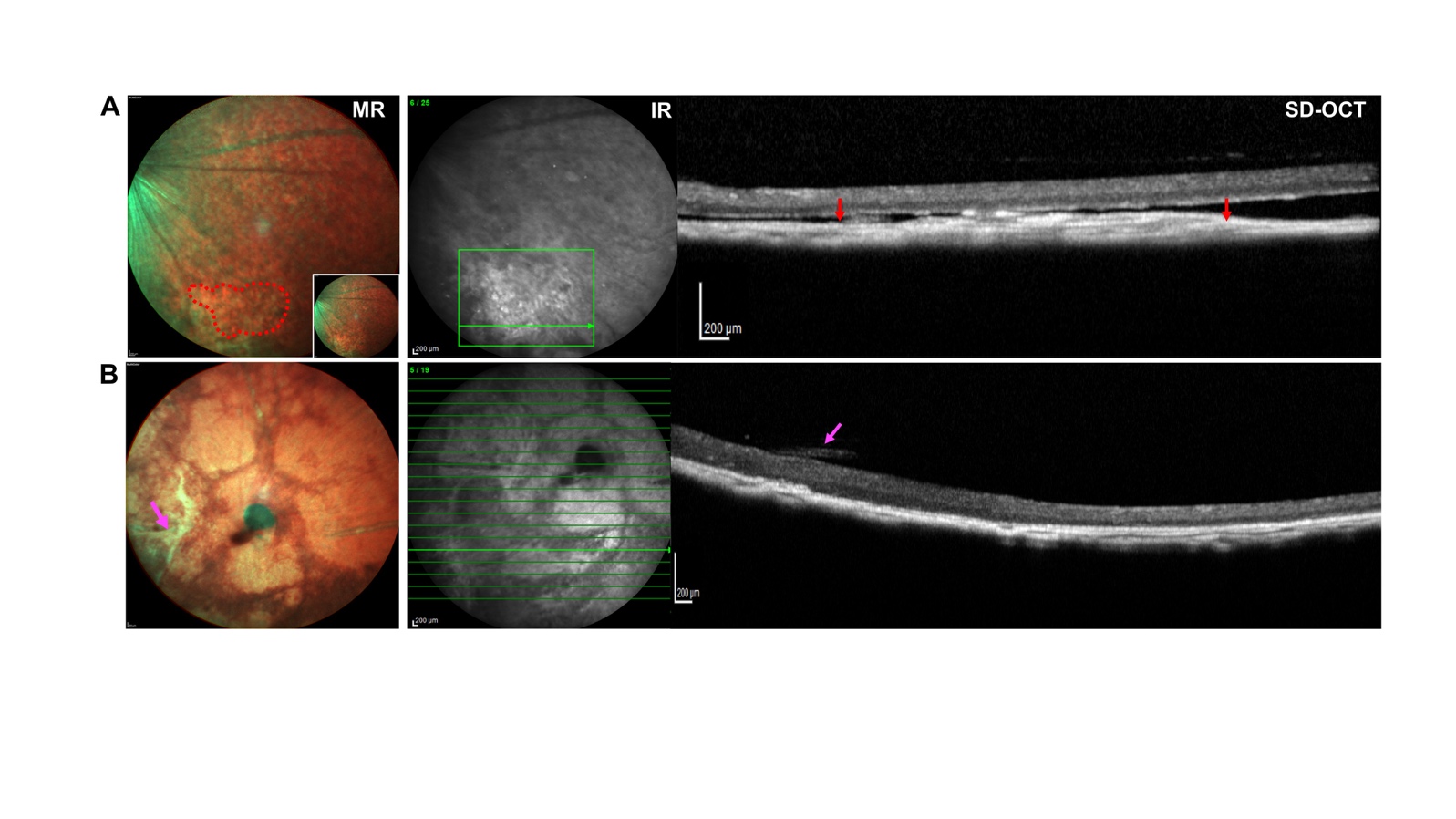

**Figure S1. Complications detected by multimodal cSLO imaging. (**A) Hemorrhage was detected as clear red puncta on MR image (within red dashed circle) (initial image is shown on the lower right corner). SD-OCT cross-section through the hemorrhage area shown on MR image showed graft-like high reflectance (between red arrows). (B) MR imaging presented a greenish structure superior to the retina. Cross-section through the same area by SD-OCT imaging found presumed peri-retinal proliferation (magenta arrow).
